# STAT1 expression in myeloid cells restrains murine norovirus-induced hepatitis and fibrosis

**DOI:** 10.64898/2026.04.26.720966

**Authors:** Ava J Sharon, Madilyn B Portas, Jasmine Wright, Harshad Ingle, Blair K Hardman, Erin J Goldberg, Jung Hee Seo, Ninan Abraham, Marc S Horwitz, Megan T Baldridge, Blayne A Sayed, Lisa C Osborne

## Abstract

**Background & Aims:** Rare cases of non-hepatotropic virus (NHV) infection in humans can cause severe hepatitis and even acute liver failure. Clinically relevant animal models of NHV-induced hepatitis are limited, contributing to the incomplete understanding of pathological mechanisms. Murine norovirus (MNV) elicits hepatosplenomegaly in mice lacking the antiviral immune effector Signal Transducer and Activator of Transcription-1 (STAT1), providing a model to investigate mechanisms of NHV-induced hepatic pathology.

**Methods:** STAT1-sufficient and -deficient (*Stat1*^Het^, *Stat1*^KO^) littermates infected intravenously (i.v.) with MNV strain CR6 were assessed for hepatic inflammation and viral burden. Cell types and molecular pathways associated with hepatic pathology in CR6-infected *Stat1*^KO^ mice were identified by flow cytometry and RNAseq of liver tissue. The relative importance of hematopoietic vs non-hematopoietic expression of STAT1 in restricting CR6 replication and maintaining tissue homeostasis was assessed in bone marrow chimeras.

**Results:** MNV CR6 *Stat1*^KO^ mice developed severe hepatitis with patchy hepatocellular necrosis and localized enrichment of CR6-infected myeloid cells, particularly macrophages. Gene set enrichment analysis (GSEA) of hepatic biopsies isolated from CR6-infected *Stat1*^KO^ mice suggested dysregulated myeloid cell activation and indicated similarities between murine and human hepatic pathologies. STAT1 expression in hematopoietic cells was protective against hepatic viral dissemination, but hematopoietic STAT1-deficiency permitted persistent hepatic MNV infection, facilitating dysregulated myeloid cell activation and hepatic fibrosis.

**Conclusions:** These results demonstrate that the role of STAT1 extends beyond restricting MNV dissemination and suggest that STAT1-dependent regulation of myeloid cell activation prevents acute hepatic necroinflammation and secondary fibrosis. This model of MNV-induced hepatitis may prove valuable in elucidating mechanisms of rare clinical complications.

**Synopsis:** Mechanisms driving acute hepatitis caused by non-hepatotropic viruses are not well understood. We describe a model of non-hepatotropic murine norovirus infection that reliably induces liver pathology and identify a requirement for STAT1 expression in myeloid cells to promote antiviral immunity and hepatic tissue protection.

**Graphical Abstract:** 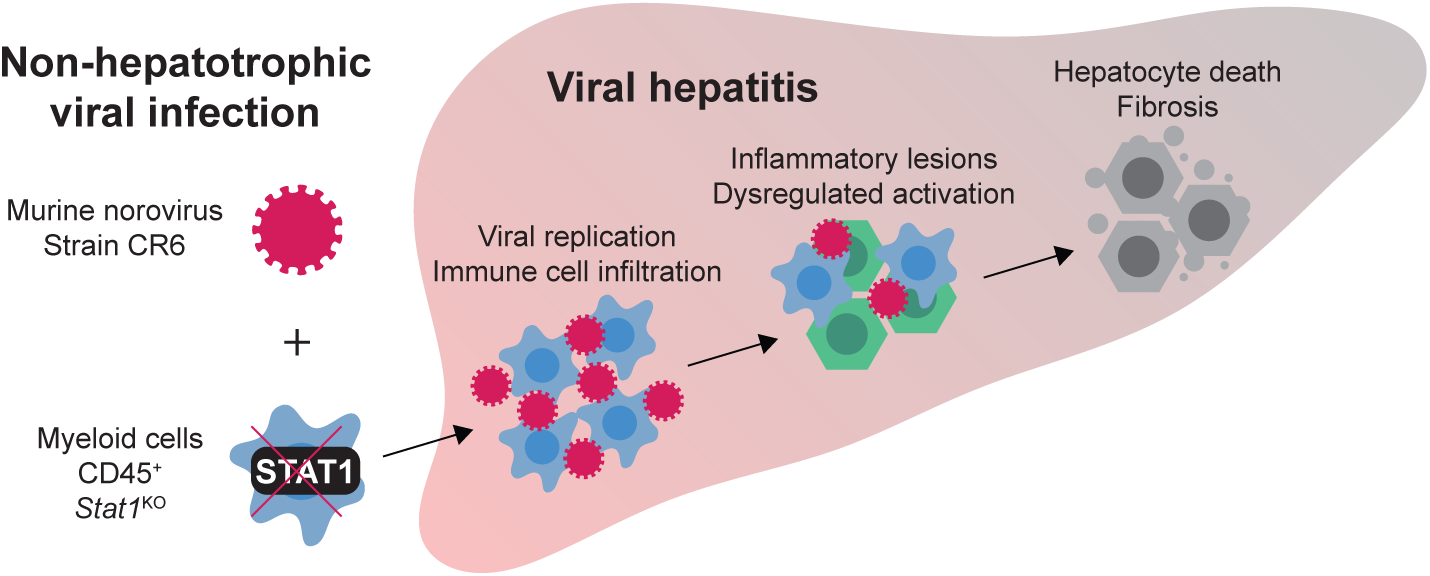

## Introduction

Severe cases of acute hepatitis, liver dysfunction, or failure secondary to non-hepatotropic viral (NHV) infections, including Epstein-Barr virus (EBV), varicella zoster, herpes simplex viruses, Parvovirus B19, rubella, measles, adenovirus and norovirus, have been reported in immunocompetent and immunocompromised adult and pediatric populations^1–9^. Notably, the etiology of ∼50% of pediatric acute liver injury cases leading to liver transplantation is unclear, but a dysregulated immune response to NHV infections has been suggested as a contributing factor^3,4,10^. In the context of genetic immunodeficiency, children with loss-of-function (LOF) mutations in Signal Transducer and Activator of Transcription-1 (STAT1) have a high risk of severe outcomes following viral infection, including Hemophagocytic Lymphohistiocytosis (HLH), an immune-mediated disease characterized by multi-organ infiltration of inflammatory myeloid cells, including the liver, and infection-induced mortality^11–19^.

Despite the severe consequences associated with NHV infections in susceptible individuals, the rarity of cases and the heterogeneity in clinical presentation contributes to our limited understanding of the mechanisms underlying NHV-mediated hepatitis. In addition, many NHVs associated with acute hepatitis have a restricted host range (humans are the only known reservoir for EBV, varicella zoster, rubella, and measles), confounding examination of pathogenesis in animal models.

Murine norovirus (MNV) is a naturally occurring murine pathogen genetically related and biologically similar to human norovirus (hNoV)^20–22^. Like many other NHV of clinical relevance, hNoV has a restricted host range, and *in vivo* approaches investigating hNoV pathogenesis are primarily limited to analysis of samples from immunocompromised patients or adapted animal models (chimpanzees, piglets, humanized mice)^23–28^. Similarities between hNoV and MNV, including genomic sequence, capsid structure, fecal-oral transmission, and the establishment of either acute or persistent infection with fecal shedding after symptomatic resolution, provided a much needed model to elucidate *in vivo* mechanisms of NoV pathogenesis^29^. A combination of advances in primary human cell culture protocols and insights from *in vivo* and *in vitro* MNV studies have revealed additional similarities between hNoV and MNV, including their ability to productively infect a variety of immune cell types (B cells, macrophage (MΦ), and other myeloid cells) and intestinal epithelial cells, interactions between capsids and bile salts, and regulation of infectious outcomes by the microbiota and host-derived factors^29–40^. Thus, a model of MNV-induced liver injury that provides insight into the immune-tissue interactions that permit and sustain hepatic inflammation could inform clinical approaches in settings of hepatitis caused by NHV infections, including hNoV.

Despite using the same cell surface receptor (CD300lf) for viral entry, two experimental strains of MNV, CW3 and CR6, demonstrate distinct cell tropism and infection dynamics in immunosufficient mice and have been used to delineate host and viral determinants of acute and persistent infectious outcomes^41,42^. MNV CW3 disseminates from the gastrointestinal tract, replicates in CD300lf-expressing myeloid lineage cells of the innate immune system (MΦ, monocytes, dendritic cells (DC), neutrophils) and B cells, and is cleared by the combined action of MNV-specific T and B cells within two weeks^43,44^. In contrast, oral inoculation with MNV strain CR6 results in persistent enteric infection. Minor genetic variants in the non-structural protein NS1 account for the ability of MNV CR6, but not CW3, to infect and persist in a rare CD300lf-expressing lineage of intestinal epithelial cells called tuft cells by evading type III interferon (IFNλ)-mediated antiviral immunity^45–47^. However, if CR6 is administered parenterally, the cellular tropism resembles that of CW3, with detectable viral burden in the spleen and scant fecal shedding^46^. This is consistent with *in vitro* findings that both MNV CW3 and CR6 readily infect and replicate in bone marrow-derived macrophages and dendritic cells (BMDM, BMDC) and MΦ-derived cell lines^21,37,48^.

Although cellular infection is influenced by interactions between MNV virions and bile salts, the liver is not significantly affected in immunocompetent hosts^21,38,39^. Early observations of vivarium acquired or deliberate MNV infection of immunodeficient mice lacking adaptive immune cells (*Rag1*^-/-^) or type I or type II interferon signaling (*Ifnar*^-/-^, *Ifngr*^-/-^) were indicative of host factors required to restrict tissue dissemination and associated tissue damage, with hepatitis as a common feature^49,50^. Multiple distinct MNV strains elicit infection-induced morbidity or mortality with high viral burdens and tissue damage in multiple organs, including the liver, in STAT1-deficient (*Stat1*^-/-^, *Stat1*^KO^) mice^51–54^. Infiltrating mononuclear phagocytes are a common feature of MNV-induced splenomegaly and hepatitis in *Stat1*^KO^ mice^22,52,53,55^, and we recently demonstrated that splenic MΦ and monocytes of CR6-infected *Stat1*^KO^ mice harbor high viral loads^51^. Despite these findings, the role of MNV infected myeloid cells and the ensuing inflammatory response in acute hepatitis and liver injury not been completely elucidated.

We recently reported that oral or systemic inoculation of MNV CR6 into *Stat1*^KO^ mice, but not their immunosufficient littermates, reliably elicited severe weight loss that was associated with hepatosplenomegaly and hepatic pathology, including foci of necrosis and inflammation^51^. In a distinct model of viral-induced hepatitis and multi-organ damage in *Stat1*^KO^ mice, CD4^+^ T cell-mediated immunopathology was responsible for the systemic inflammation and mortality caused by lymphocytic choriomeningitis virus (LCMV)^56^. Despite a similar hyperaccumulation of pro-inflammatory CD4^+^ and CD8^+^ T cells in tissues of CR6-infected *Stat1*^KO^ mice, adaptive immune cell depletion (CD4^+^ or CD8^+^ T cells alone, or CD4/CD8 T cells and CD20^+^ B cells together) had no effect on morbidity or mortality^51^. Instead, tissue and clinical protection was achieved by therapeutic treatment with an antiviral that restricted, but did not eliminate, systemic CR6 infection in *Stat1*^KO^ mice. These data suggest an alternative mechanism of STAT1-dependent protection, but the STAT1-expressing cell types that restrict hepatic CR6 dissemination and prevent tissue pathology have not been defined. In particular, whether STAT1-deficient non-hematopoietic hepatic cells become permissive for CR6 replication or contribute to tissue injury in other ways is unknown.

Here, we use systemic infection of *Stat1*^KO^ with MNV strain CR6 to examine mechanisms of NHV-induced induced hepatitis and collateral tissue damage that may have relevance for clinical cases of liver injury secondary to hNoV and other NHV infections. We demonstrate the emergence of focal hepatic lesions enriched with myeloid cells co-expressing MNV proteins in CR6-infected *Stat1*^KO^ mice, suggesting *in vivo* permissiveness for CR6 replication. Transcriptome analysis revealed signs of myeloid cell dysregulation, including expression of genes associated with alternative activation, collagen deposition, and fibrosis in the CR6-infected STAT1-deficient liver. Consistent with the implication that STAT1-deficient myeloid cells establish and propagate NHV pathogenesis, bone marrow chimeras demonstrated that STAT1 expression in leukocytes prevented CR6-induced viral dissemination and hepatic pathology. In contrast, STAT1-deficiency in the hematopoietic compartment resulted in persistent hepatic infection, fibrosis, and signs of alternatively activated macrophages (AAMΦ). These findings demonstrate that STAT1-dependent regulation of innate immune cells prevents acute MNV dissemination, and elucidate a critical role for STAT1 in immune-mediated control of post-viral sequelae.

## Results

### Localized areas of liver pathology and MNV CR6 replication in STAT1-deficient mice

We previously reported that oral infection with MNV CR6 led to ∼50% of infected *Stat1*^KO^ mice developing infection-induced morbidity by 7 days post-infection (pi), whereas systemic infection by the intravenous (i.v.) route led to reliable CR6-induced disease that met the requirements for humane endpoint by day 7 pi^51^. Notably, systemic CR6 infection was asymptomatic in *Stat1*^Het^ littermates^51^. Systemic CR6-induced disease in *Stat1*^KO^ mice is characterized by severe weight loss and multi-organ tissue pathology, including hepatosplenomegaly with gross evidence of numerous pale nodules distributed throughout the organs (Fig. 1A-C), phenotypes that are consistent with reports from independent investigators across multiple facilities^10–12^. Quantification of hepatic tissue damage in sections of the left lateral lobe at day 7 post-CR6 infection supported prior results indicating that STAT1 expression restrains CR6 replication and prevents hepatic damage, and that impaired control of viral replication in *Stat1*^KO^ littermates is associated with focal patches of hepatic inflammation and non-confluent necrosis distributed across >10% of the area examined in liver sections (Fig. 1D-F). Together, these data provide the foundation for using this as a model of immunodeficient NHV pathogenesis.

**Figure 1:**
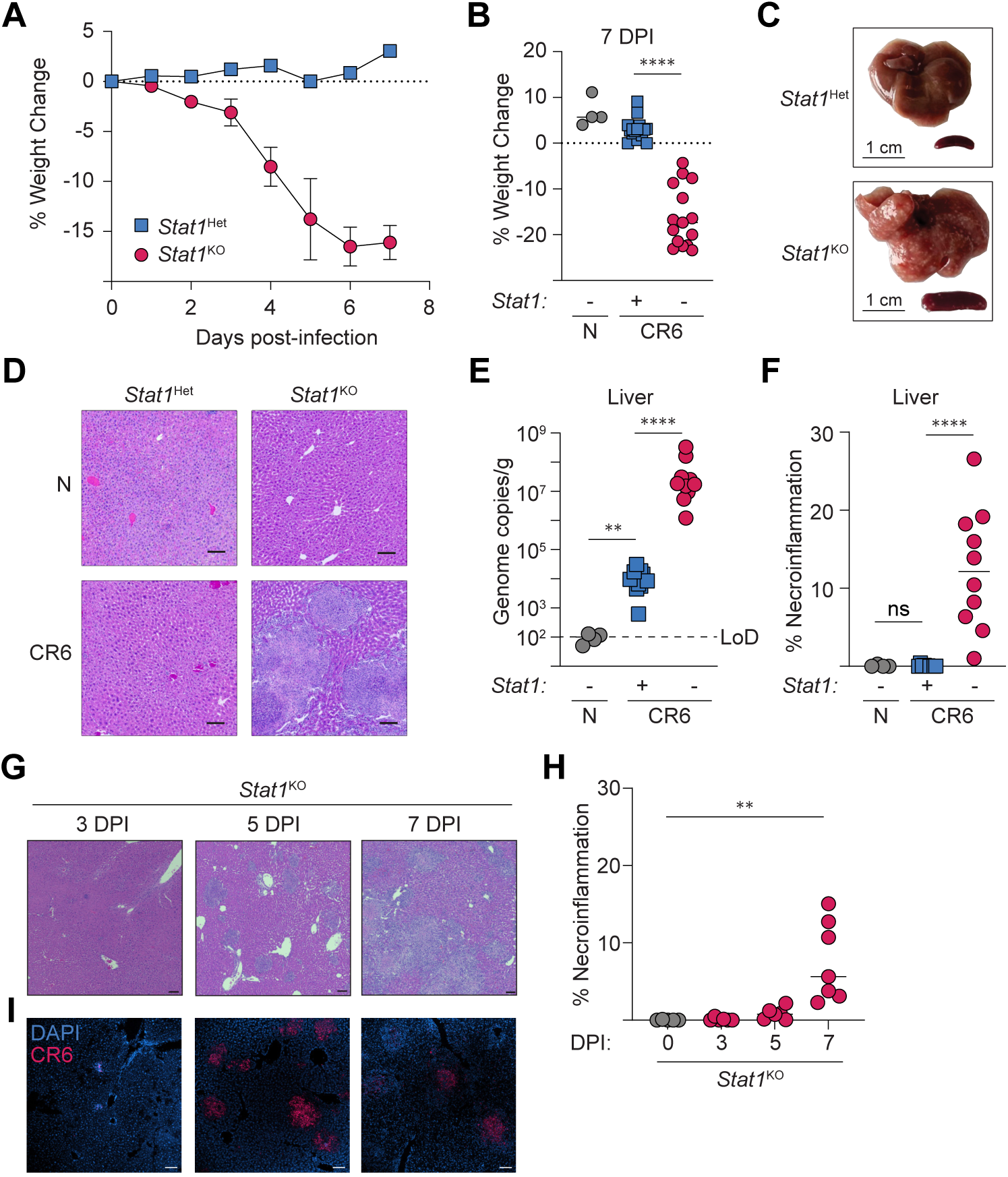
Localized areas of liver pathology and MNV CR6 replication in STAT1-deficient mice. *Stat1*^Het^ and *Stat1*^KO^ littermates were i.v. infected with 2x10^5^ PFU of CR6 (or sham-infected) and analyzed at 7 days pi (A-F) or indicated timepoints (G-I). (A, B) Percent weight change, n=4 naive and n=15-16 MNV CR6-infected *Stat1*^Het^ and *Stat1*^KO^. (C) Whole liver and spleen at d7 pi. (D) Representative images of H&E-stained, FFPE liver sections. (E) Hepatic MNV CR6 genome copies detected by qRT-PCR, normalized to tissue weight. (F) Quantified areas of necroinflammation in liver sections. (D-F) n=4 naive and n=10 MNV CR6-infected *Stat1*^Het^ and *Stat1*^KO^. (G, H) Representative images (G) and quantified areas of necroinflammation (H) in H&E-stained, FFPE liver sections of naive and CR6-infected *Stat1*^KO^ (n=5-8 per timepoint). (I) RNAscope detection of MNV CR6 genomes and DNA (DAPI). All graphs are pooled from at least two independent experiments with n=4 naive and n=15-16 MNV CR6-infected *Stat1*^Het^ and *Stat1*^KO^. Images are representative of two independent experiments. Mann-Whitney U test with Holm-Šídák correction for multiple comparisons where applicable. ** = p < 0.01, **** = p < 0.0001, ns = not significant. Scale bars = 100 µm unless otherwise indicated. LoD = limit of detection.

We hypothesized that hepatic lesions could be established by or include the initial waves of MNV CR6-infected cells. To assess this, we performed a kinetic analysis of CR6-induced hepatic pathology, demonstrating the presence of focal inflammatory infiltrates at day 3, with continued expansion at day 5 that culminated in enlarged regions of necroinflammation by day 7 pi (Fig. 1G, H). RNAScope imaging revealed the presence of CR6 genomic material in cells with myeloid morphology at day 3 pi and in tight clusters that matched the distribution of focal liver lesions at day 5 pi (Fig. 1I). Notably, nuclear DAPI staining was diffuse in areas surrounding viral genome clusters in *Stat1*^KO^ livers at days 5 and 7 pi, suggesting that CR6-induced cell death drives hepatocellular necrosis (Fig. 1I). Together, these data suggest that impaired antiviral immunity in CR6-permissive cells may initiate the development and spread of focal areas of unrestrained viral replication and necroinflammation.

### STAT1-deficient myeloid cells infiltrate the liver and are permissive for MNV CR6 replication

Our histological findings indicated a correlation between areas of hepatic necrosis and sites of CR6-infected cells in *Stat1*^KO^ mice, but they did not identify the cell type(s) hosting viral replication. To address this, we quantified viral genomes in flow-sorted populations of CD45^+^ leukocytes and CD45^-^ non-hematopoietic cells isolated from *Stat1*^Het^ and *Stat1*^KO^ littermates at day 5 post-CR6 infection. CR6 was not detected in cells sorted from STAT1-sufficient animals but viral genome copies were present in both CD45^-^ and CD45^+^ populations isolated from *Stat1*^KO^ mice, albeit enriched by ∼100-fold in CD45^+^ cells (Fig. 2A).

**Figure 2:**
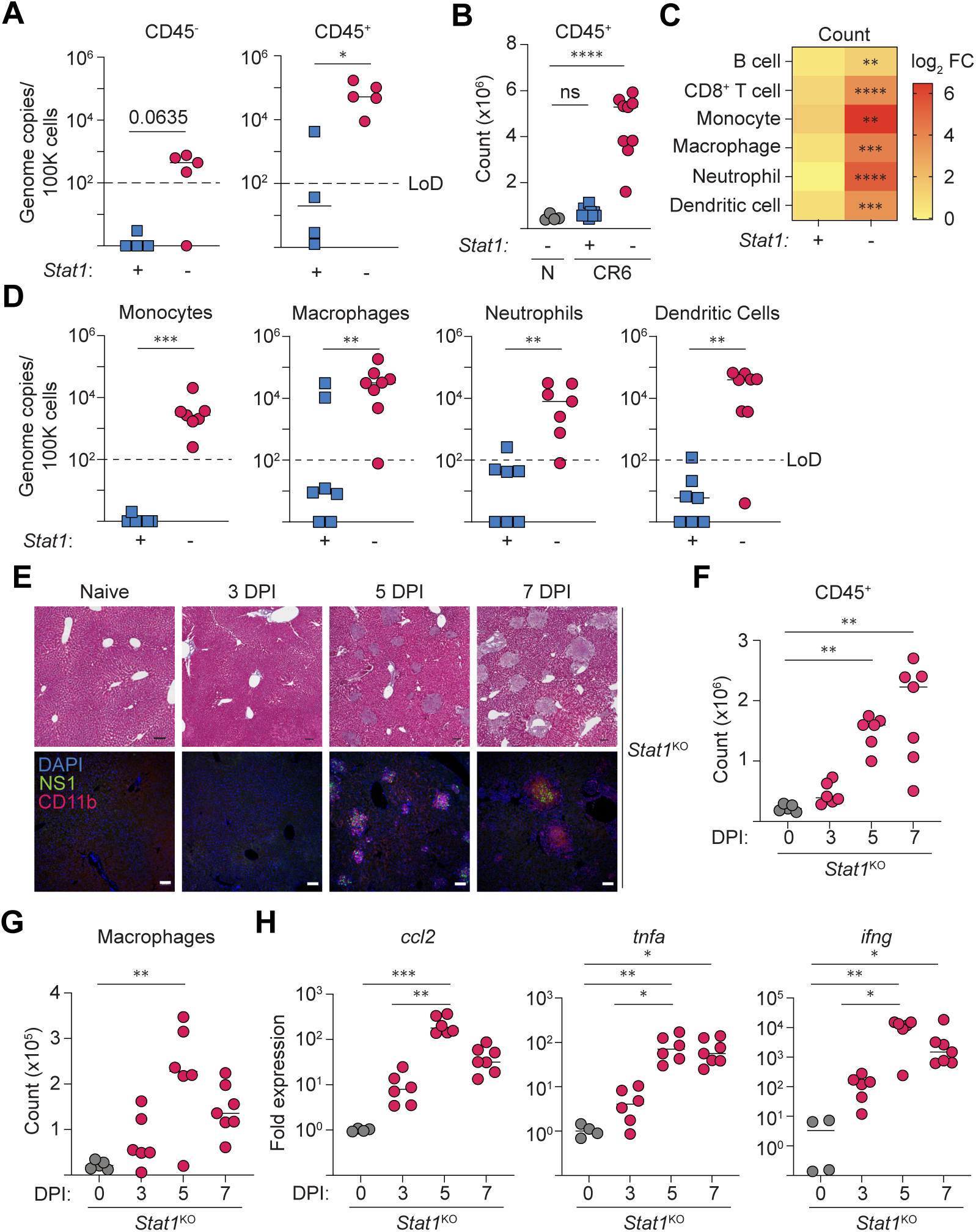
STAT1-deficient myeloid cells infiltrate the liver and are permissive for MNV CR6 replication. *Stat1*^Het^ and *Stat1*^KO^ mice were i.v. infected with 2x10^5^ PFU of CR6 (or sham-infected) and analyzed at 5 days pi (A-D) or the times indicated (E-H). All flow-based measurements represent cells recovered from the left lateral lobe of the liver. (A) Quantification of CR6 genome copies in flow-sorted CD45^-^ and CD45^+^ cells. n=4-5 per group from 1 independent trial. (B) Quantification of live CD45^+^ leukocytes isolated from the liver left lateral lobe. (C) Total numbers of hepatic CD45^+^ immune populations, represented in the heatmap as the log_2-_fold change relative to uninfected controls (CD19^+^ TCRβ^-^ B cells, CD19^-^ TCRβ^+^ CD8α^+^ T cells, myeloid lineage cells all pre-gated on CD19^-^ TCRβ^-:^ CD11b^+^ F4/80^-^ CD11c^-^ Ly6C^+^ Ly6G^-^ monocytes; CD11b^+^ F4/80^+^ MΦ, CD11b^+^ F4/80^-^ CD11c^-^ Ly6C^+^ Ly6G^+^ neutrophils, F4/80^-^ CD11c^+^ DCs). (D) RT-qPCR quantification of CR6 genome copies in flow-sorted cell populations (gating strategy as in C). Data in B-D are compiled from 2 independent experiments with n=4 naive, n=7 *Stat1*^Het^, and n=10 *Stat1*^KO^ CR6-infected mice. (E) Naive or CR6-infected *Stat1*^KO^ FFPE liver sections stained for CD11b, MNV NS1, and DNA (DAPI) (top) or Masson’s Trichrome (bottom). Scale bars = 100 µm. Representative images from 2 independent experiments. (F, G) Quantification of live CD45^+^ leukocytes, and CD45^+^ CD19^-^ TCRβ^-^ CD11b^+^ F4/80^+^ MΦ recovered from the left lateral lobe of naive or CR6-infected *Stat1*^KO^ livers. (H) Gene expression of *ccl2, tnfa,* and *ifng* in liver RNA, normalized to *hprt* expression, expressed as fold-change compared to uninfected controls. Data are pooled from two independent experiments, n=5-7 per group. Mann-Whitney U test, with Holm-Šídák correction for multiple comparisons where applicable (B, F, H). * = p < 0.05, ** = p < 0.01, *** = p < 0.001, **** = p < 0.0001, ns = not significant. Scale bars = 100 µm. LoD = limit of detection.

Compared to naïve counterparts (*Stat1*^Het^ and *Stat1*^KO^) and CR6-infected *Stat1*^Het^ mice, the livers of CR6-infected *Stat1*^KO^ mice contained an expanded population of infiltrating CD45^+^ leukocytes at day 5 pi (Fig. 2B) that were candidates for viral replication, including CD19^+^ B cells, CD8^+^ T cells, and myeloid lineage monocytes, MΦ, neutrophils, and DCs (Fig. 2C). Although splenic T and B cells can harbor MNV CR6, they are dispensable for CR6-induced morbidity in *Stat1*^KO^ mice^51^. Thus, we examined hepatic myeloid cell populations.

Consistent with reports that *Stat1* immunodeficiency leads to broad cellular tropism in MNV CW3-infected *Stat1*^KO^ mice^36^, CR6 genome copies were enriched in multiple myeloid-lineage cells isolated from the livers of *Stat1*^KO^ mice, while viral loads in STAT1-sufficient control cells were at or below the limit of detection (except for 2 samples with detectable levels in isolated hepatic MΦ) (Fig. 2D). Further, immunostaining of MNV non-structural protein 1 (NS1) as a proxy for viral infection demonstrated foci of co-localized MNV NS1- and CD11b-expression by day 5 pi. Notably, despite qPCR-mediated detection of MNV genomes in flow-sorted CD45^-^ cells, by day 7 pi, MNV NS1 staining was largely restricted to the areas of non-confluent necrosis with clustered CD11b^+^ cells in CR6-infected *Stat1*^KO^ mice (Fig. 2E). Kinetic analyses demonstrated that total MΦ cell numbers began to increase at day 3 and remained elevated through 7 days of CR6 infection in *Stat1*^KO^ mice. Notably, MΦ recruitment coincided with elevated hepatic expression of genes coding for the monocyte chemoattractant *ccl2* and inflammatory cytokines *ifng* and *tnfa* (Fig. 2F-H). Collectively, these data suggest a STAT1-dependent mechanism of tissue protection that, when disrupted, results in the accumulation of MNV-infected myeloid cells that precipitate hepatic necroinflammation.

### The transcriptional landscape of CR6-infected *Stat1*^-/-^ livers resembles other hepatopathologies and indicates disrupted metabolism and myeloid cell activation

To broaden our understanding of the consequences of CR6-induced hepatitis, we performed RNA sequencing on liver biopsies from the left lateral lobe of *Stat1*^KO^ and *Stat1*^Het^ mice at day 5 post-CR6 infection. More than 800 genes were differentially expressed (adjusted p-value < 0.05 and log_2_FC > 2), including up- and down-regulation of 581 and 282 genes, respectively, in *Stat1*^KO^ compared to *Stat1*^Het^ littermates (Fig. 3A). Gene set enrichment analysis (GSEA) comparison to publicly available datasets indicated significant similarities between the transcriptional signatures of CR6-infected *Stat1*^KO^ livers and non-infectious liver pathologies of wild-type (WT) mice, including autoimmune hepatitis^57^, IL-1β injection^58^, acute alcohol toxicity^59^, and hepatic fibrosis^60^ (Fig. 3B). We also annotated gene sets derived from liver biopsies of patients experiencing acute Hepatitis B-induced liver failure to facilitate comparison to murine homologs^61^. Providing further support for the translational relevance of this model, GSEA revealed significant overlap in the transcriptional landscape of livers from Hepatitis B patients and CR6-infected *Stat1*^KO^ mice (Fig. 3C).

**Figure 3:**
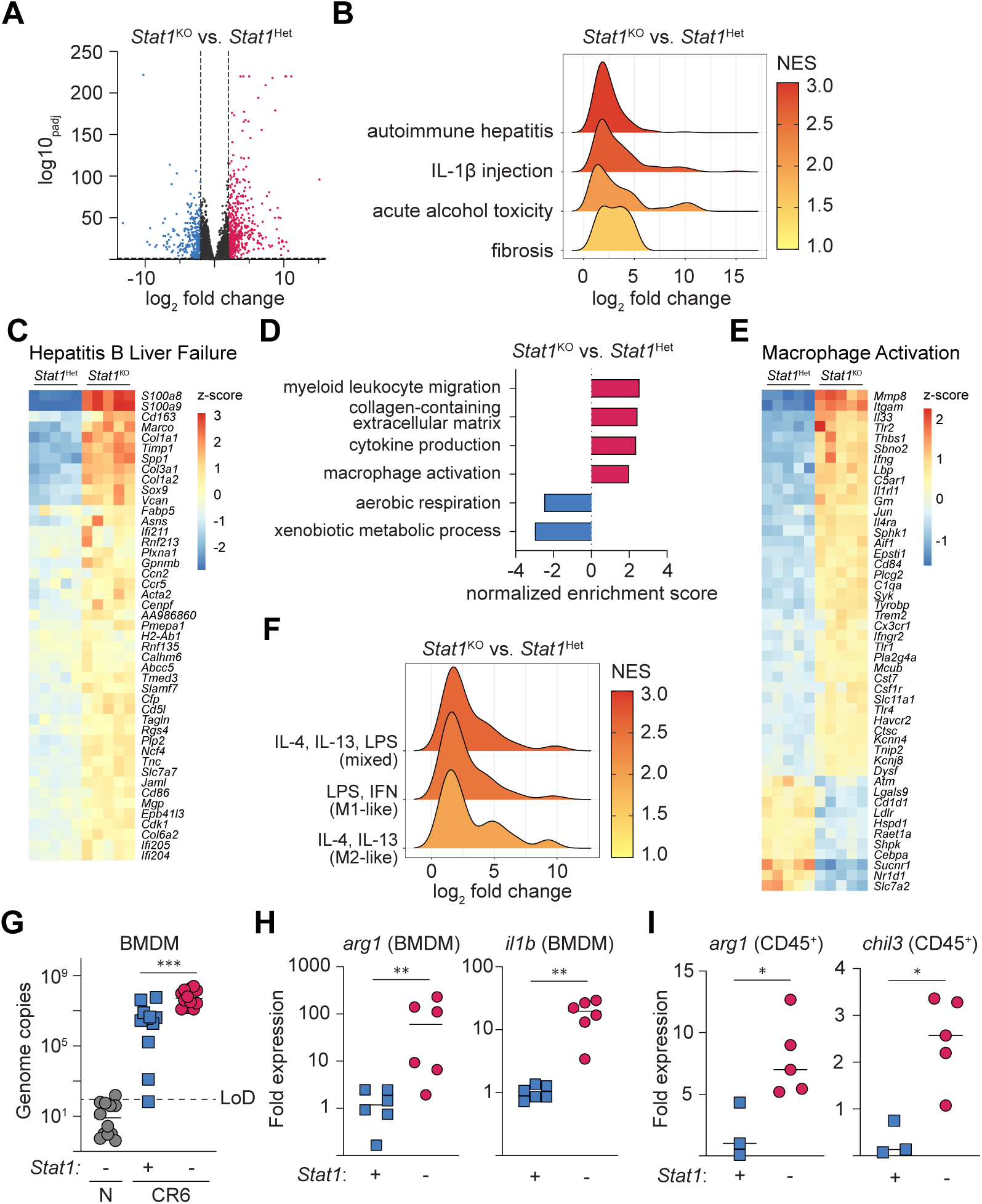
Liver disease and macrophage-associated pathways are activated in the liver during CR6-induced disease. *Stat1*^Het^ and *Stat1*^KO^ mice were i.v. infected with 2x10^5^ PFU of CR6 (n = 5 mice per group). At 5 days pi, RNA was extracted from the left lateral lobe of the liver for sequencing (A-F). (A) Volcano plot of all analyzed genes. Data points are coloured if p_adj_ < 0.05 and log_2_ fold change > or < 2. (B) GSEA of published liver disease gene sets (autoimmune hepatitis: GSE9892, IL-1β injection: GSE216278, alcohol toxicity: GSE214778, and fibrosis: GSE222576) significantly enriched in *Stat1*^KO^ mice. (C) Genes from a published analysis of human acute Hepatitis B liver failure (GSE62037) were converted to mouse IDs (see methods). The heatmap represents genes enriched in both acute Hepatitis B liver failure and CR6-infected *Stat1*^KO^ mice (NES = 2.99, padj < 0.00001). (D) GSEA was performed against Gene Ontology Biological Process terms. Selected significant terms differentially up- or down-regulated in *Stat1*^KO^ mice are shown. (E) Heatmap of the *Macrophage Activation* GO term (NES = 1.90, p_adj_ = 0.0002) arranged by unsupervised hierarchical clustering. (F) GSEA comparison between CR6-infected *Stat1*^KO^ liver tissue and published gene sets from BMDM activated by LPS & IFNγ, IL-4 & IL-13, or both (GSE138263). (G-H) *Stat1*^Het^ and *Stat1*^KO^ BMDM were infected with 5x10^5^ PFU of CR6, and RNA was extracted 24 hours pi. (G) RT-qPCR quantification of CR6 genome copies in cultured BMDM. (H) Gene expression of *il1b* and *arg1* in CR6-infected BMDM, normalized to *hprt* expression, and expressed as fold-change relative to uninfected controls. (I) Relative expression of AAMΦ signature genes *arg1* and *chil3* in flow-sorted CD45^+^ cells isolated from the left lateral lobe of CR6-infected *Stat1*^Het^ and *Stat1*^KO^ mice at 5 days pi, normalized to *hprt* expression and uninfected controls. Data are pooled from two (H), three (G), or one (I) independent experiments. Mann-Whitney U-test. ** = p < 0.01, *** = p < 0.001. NES = normalized enrichment score. LoD = limit of detection.

Gene Ontology (GO) biological process term analysis suggested an overall disruption of liver function in CR6-infected *Stat1*^KO^ mice, characterized by diminished expression of genes associated with aerobic respiration and xenobiotic metabolic processes (Fig. 3D). GO term analysis also indicated enrichment of extracellular matrix remodeling, consistent with similarities to hepatic fibrosis, and a complex inflammatory milieu, including significant enrichment of signatures associated with myeloid cell migration, cytokine production, and MΦ activation (Fig. 3D). These observations were supported by robust differences in gene expression related to MΦ activation (Fig. 3E). Notably, genes associated with both classical (type I and type II IFNs) and alternative MΦ activation (*il4, il13*) were enriched in the livers of CR6-infected *Stat1*^KO^ hosts (Fig. 3F), consistent with previous studies indicating a requirement for STAT1 to guide appropriate MΦ polarization^62,63^.

Transcriptome analysis was performed on bulk liver biopsies. Thus, to examine the consequences of CR6 infection on MΦ polarization, we generated and infected *Stat1*^Het^ and *Stat1*^KO^ BMDM. At 24 hrs pi, MNV had replicated in BMDM of both genotypes, albeit with a more uniform distribution in *Stat1*^KO^ cells (Fig. 3G). Alongside up-regulation of genes associated with classical MΦ activation (e.g, *il1b*), genes associated with AAMΦ (*arg1)* were also enriched in infected *Stat1*^KO^ BMDM (Fig. 3H). Consistent with previous studies indicating a requirement for STAT1 to guide appropriate MΦ polarization^62–64^, AAMΦ signature genes *arg1* and *chil3* were elevated in sorted CD45^+^ cells from CR6-infected *Stat1*^KO^ mice (Fig. 3I). Notably, products of AAMΦ are associated with extracellular remodelling and fibrosis^65,66^. Together, these data provide new insights into the hepatic pathology caused by CR6 infection of *Stat1*^KO^ mice. In addition to suggesting altered hepatocyte metabolism (potentially due to cell death), these data indicate that dysregulated myeloid cell activation could subsequently contribute to NHV-induced hepatic inflammation and fibrosis in STAT1-deficient mice.

### STAT1 expression in hematopoietic cells is necessary and sufficient for protection from CR6-induced disease

Taken together, our data indicate that liver pathology is associated with general liver metabolic dysfunction and the infiltration of CR6-permissive myeloid cells that fail to contain viral replication and undergo dysregulated activation. However, these associations are insufficient to define the STAT1-expressing cell types that coordinate tissue protection following non-hepatotropic MNV infection. To investigate this, we generated bone marrow (BM) chimeric animals with congenic CD45.2^+^ *Stat1*^KO^ mice and wild-type (WT) CD45.1^+^ BoyJ mice to restrict STAT1 expression to either hematopoietic stem cell (HSC)-derived donor leukocytes or host-derived radioresistant cells, including hepatocytes (Fig. 4A). At 6 weeks post-reconstitution, >75% of the hematopoietic compartment was derived from congenic bone marrow donors, and mice were infected with CR6 (Fig. 4B).

**Figure 4:**
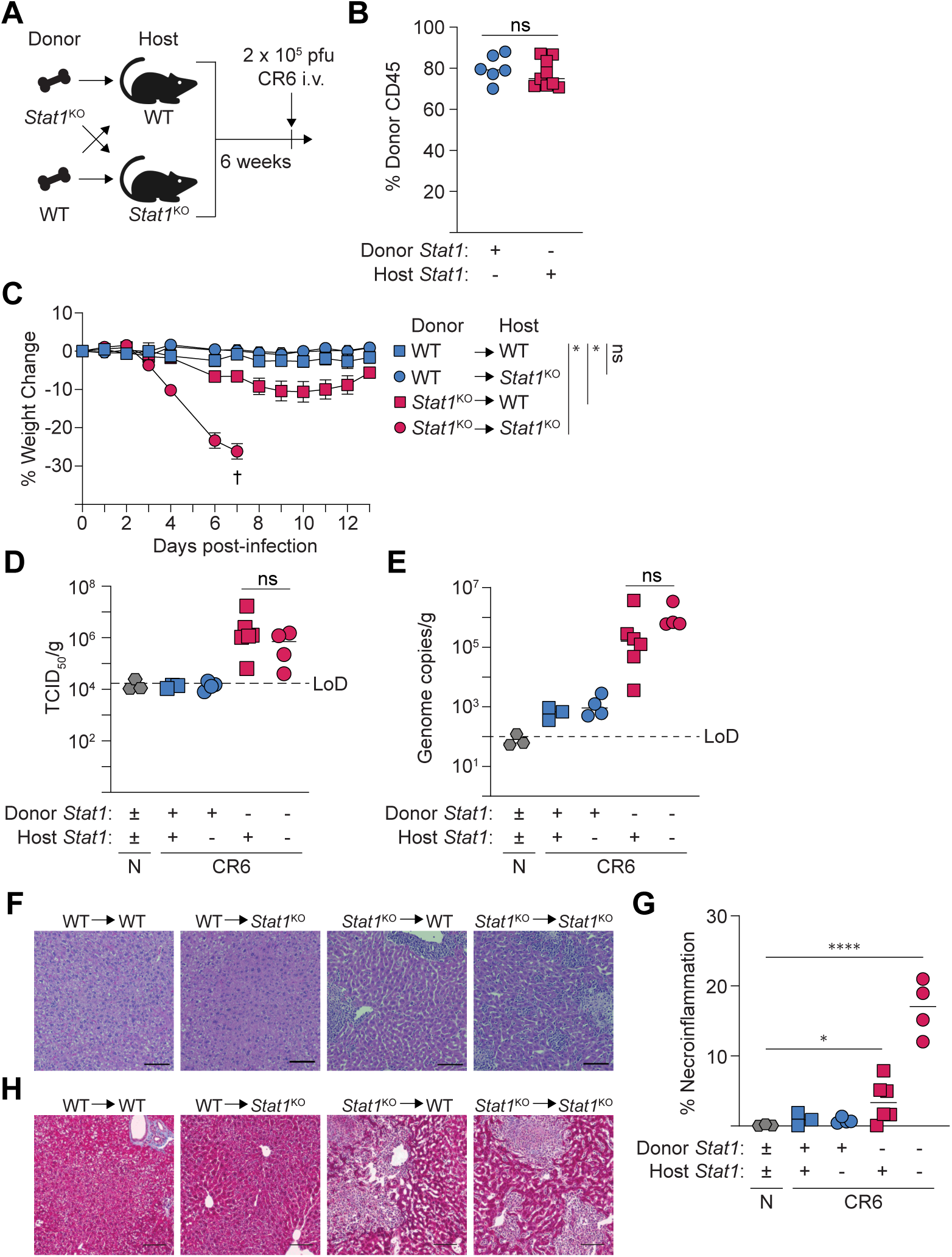
**STAT1 expression in hematopoietic cells is necessary and sufficient for protection from CR6-induced liver disease**. (A) Experimental design for bone marrow chimera experiments. Lethally irradiated *Stat1*^KO^ (CD45.2^+^) and WT (BoyJ, CD45.1^+^) mice received i.v. bone marrow transplants from sex-matched *Stat1*^KO^ or WT donors. After 6 weeks of reconstitution, mice were infected with 2x10^5^ PFU of CR6 and analyzed at day 7 pi. (B) Percent expression of donor congenic CD45 marker on PBMCs before infection. (C) Percent weight change from initial over time. † indicates euthanasia due to humane endpoint. (D) TCID_50_ quantification of infectious CR6 in liver lysate normalized to tissue weight. (E) Quantification of hepatic CR6 genome copies, normalized to tissue weight. (F, G) H&E staining (F) and (G) quantified areas of necroinflammation in FFPE liver sections of CR6-infected mice. (H) Masson’s trichrome staining of FFPE livers at 7 days post CR6-infection. Scale bars = 100 µm. Data are pooled from two independent experiments (B-C, n=7-17 per group), or are representative of two independent experiments (D-G). For (C), the extra sum-of-squares F test was performed on fifth-order polynomial lines fitted to both groups or to each group individually. For all other significance testing, Mann-Whitney U-test with Holm-Šídák correction for multiple comparisons. * = p < 0.05, **** = p < 0.0001, ns = not significant. LoD = limit of detection.

Following CR6 infection, mice with a STAT1-sufficient hematopoietic compartment showed no clinical signs of disease. Independent of host genotype, WT → WT and WT → *Stat1*^KO^ BM chimeras maintained body weight and effectively prevented hepatic CR6 replication in the liver, (Fig. 4C-E). Like unmanipulated STAT1-deficient mice, CR6 infection caused rapid and severe (≥20%) weight loss in *Stat1*^KO^ → *Stat1*^KO^ chimeras that required humane intervention (euthanasia) by day 7 pi (Fig. 4C). In contrast to all other groups, *Stat1*^KO^ → WT chimeras exhibited milder (∼10%) and transient weight loss that was nearly fully recovered by day 14 pi (Fig. 4C). Despite the reduced susceptibility to infection-induced weight loss and morbidity in *Stat1*^KO^ → WT chimeras, hepatic viral loads (CR6 infectious viral burden and genome copies) were indistinguishable from *Stat1*^KO^ → *Stat1*^KO^ chimeras at day 7 pi (Fig. 4D, E). These data demonstrate that STAT1 expression in hematopoietic-derived immune cells is both necessary and sufficient to restrict CR6 dissemination to the liver.

Consistent with effective prevention of hepatic infection, there were no signs of liver injury in mice with a STAT1-sufficient hematopoietic compartment (Fig. 4F,G). In contrast, chimeras reconstituted with *Stat1*^KO^ BM developed characteristic signs of CR6-induced hepatic pathology, including foci of necroinflammation (Fig. 4F). However, the total affected area was diminished in *Stat1*^KO^ → WT compared to *Stat1*^KO^ → *Stat1*^KO^ chimeras (Fig. 4F, G). Finally, we reasoned that the enriched transcriptomic signatures of collagen-containing extracellular matrix and AAMΦ (Fig. 3D-F) may suggest that the livers of CR6-infected *Stat1*^KO^ mice were susceptible to fibrosis. We assessed collagen deposition by Masson’s trichrome staining in the livers of BM chimeras at day 7 pi. Some indication of collagen deposition was detected in hepatic lesions of *Stat1*^KO^ → WT and *Stat1*^KO^ → *Stat1*^KO^ chimeras, but overt fibrosis was not apparent at this timepoint (Fig. 4H).

Collectively, these data support a critical role for STAT1-mediated antiviral immunity in restraining CR6 replication in permissive leukocytes and preventing hepatic dissemination. Importantly, they also demonstrate that despite robust hepatic viral replication in STAT1-deficient immune cells, severe infectious outcomes were mitigated in *Stat1*^KO^ → WT BM chimeras, thereby implicating a requirement for STAT1 expression in non-hematopoietic-derived cells to maintain tissue homeostasis in the context of viral infection.

### Hematopoietic STAT1 deficiency permits persistent hepatic CR6 infection, macrophage dysregulation, and fibrosis

The survival advantage of CR6-infected *Stat1*^KO^ → WT chimeras compared to *Stat1*^KO^ → *Stat1*^KO^ cohorts provided an opportunity to analyze longer-term consequences of STAT1 immunodeficiency on post-infection sequelae. To this end, we assessed hepatic viral loads, immune infiltration, and tissue histopathology in the cohorts of chimeras that survived CR6 infection.

Independent of the genotype of the host, chimeras reconstituted with a STAT1-sufficient hematopoietic compartment (WT → WT or WT → *Stat1*^KO^) had no evidence of hepatic CR6 infection (Fig. 5A). Further, although the total number of recruited immune cells was limited in chimeras with WT BM, the primary cell types detected were CD8^+^ T cells and MΦ (Fig. 5B). Consistent with the idea that hematopoietic-derived STAT1 expression prevents CR6 dissemination and protects host tissue, neither hepatitis or fibrosis were detected in H&E or Masson’s Trichrome stained liver sections of these mice (Fig. 5C, D).

**Figure 5:**
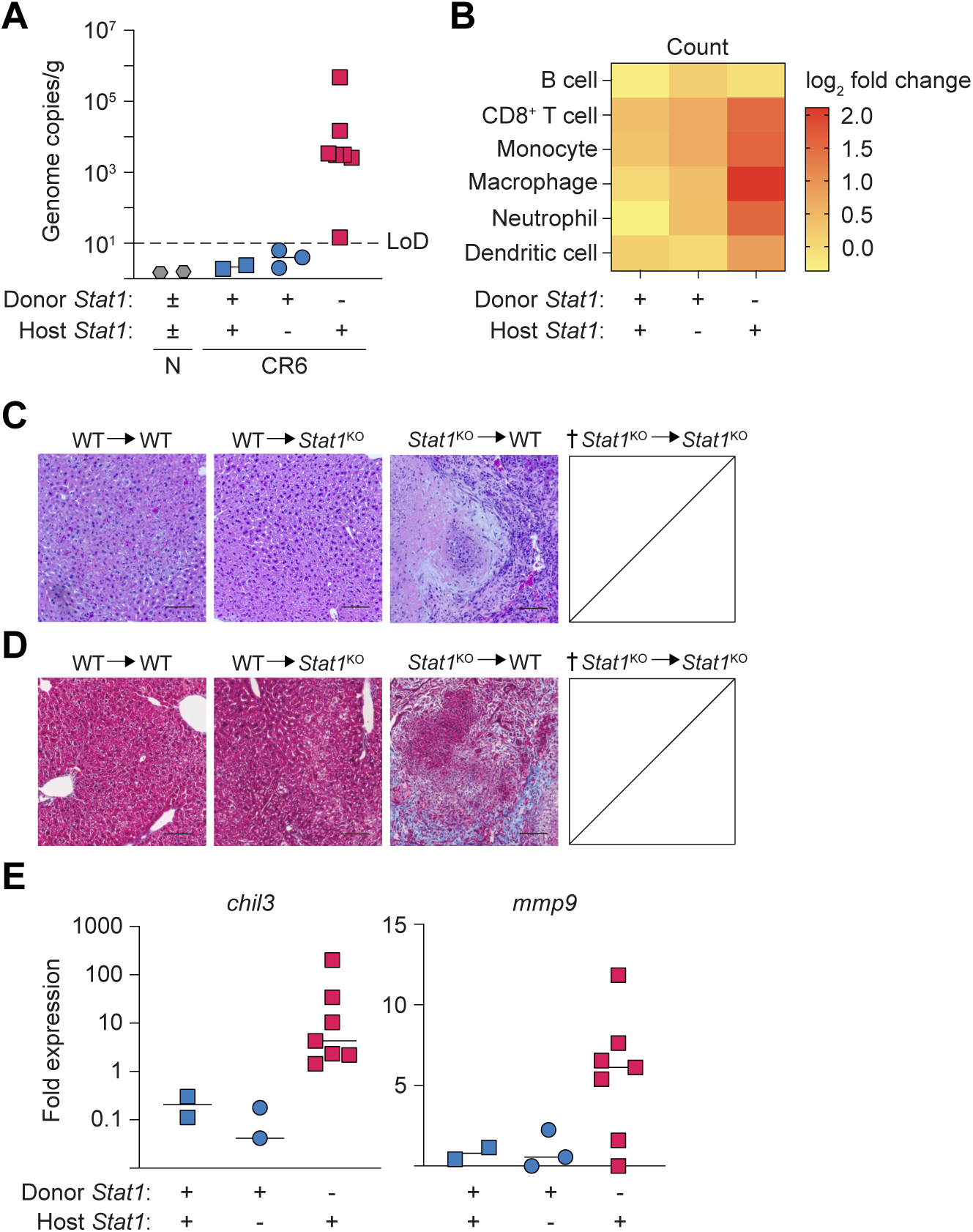
Hematopoietic STAT1 deficiency permits persistent hepatic CR6 infection, macrophage dysregulation and fibrosis. Bone marrow chimeras were generated as in Figure 4 and analyzed at day 16 post-CR6 infection. *Stat1*^KO^ → *Stat1*^KO^ chimeras reached humane endpoint at day 7 pi, precluding analysis at the later timepoint. (A) Quantification of hepatic CR6 genome copies, normalized to tissue weight. (B) Total numbers of CD45^+^ immune populations isolated from the left lateral lobe of the liver, represented in the heatmap as the log_2-_fold change relative to uninfected controls (CD19^+^ TCRβ^-^ B cells, CD19^-^ TCRβ^+^ CD8α^+^ T cells, myeloid lineage cells all pre-gated on CD19^-^ TCRβ^-^ non-B/non-T cells: CD11b^+^ F4/80^-^ CD11c^-^ Ly6C^+^ Ly6G^-^monocytes; CD11b^+^ F4/80^+^ MΦ, CD11b^+^ F4/80^-^ CD11c^-^ Ly6C^+^ Ly6G^+^ neutrophils, F4/80^-^ CD11c^+^ dendritic cells). (C, D) H&E or Masson’s trichrome staining of FFPE livers at day 16 pi. (E) Expression of fibrosis-related genes *chil3* and *mmp9* in the liver of CR6-infected chimeras, normalized to *hprt* expression and expressed as fold-change relative to uninfected controls. Scale bars = 100 µm. Data are representative of two independent experiments with n=2-7 mice per group. LoD = limit of detection.

In contrast, despite the clinical recovery, surviving *Stat1*^KO^ → WT chimeras continued to harbor hepatic CR6 infection (Fig. 5A), although the absolute magnitude of infection had reduced by ∼10^2^-fold in liver tissue collected at day 16 compared to day 7 pi. Consistent with our prior observations^51^, the hepatic CD8^+^ T cell population was ∼2-fold higher in *Stat1*^KO^ → WT chimeras compared to chimeras with WT BM (Fig. 5B). Notably, MΦ were the most abundant leukocyte population in the livers of *Stat1*^KO^ → WT chimeras (Fig. 5B), which was associated with enriched expression of genes associated with AAMΦ (*chil3*) and extracellular matrix remodelling and fibrosis (*mmp9*) (Fig. 5E). Further, expression of *chil3* correlated with viral load in the livers of *Stat1*^KO^ → WT chimeras (Spearman’s correlation coefficient: r^2^ = 0.9917, p < 0.0001), suggesting a potential infectious dose-dependent regulation of MΦ polarization. Consistent with our hypothesis that AAMΦ could render CR6-infected tissues susceptible to fibrosis, Masson’s trichrome staining revealed large, diffuse regions of collagen deposition throughout liver sections of clinically recovered *Stat1*^KO^ → WT chimeras (Fig. 5C, D).

Collectively, these data indicate that STAT1-mediated antiviral immunity in hematopoietic cells is required not only to prevent persistent hepatic MNV infection and acute hepatitis, but that the associated dysregulation of myeloid cell activation and polarization contributes to collateral tissue damage and fibrosis.

## Discussion

Here, we report initial findings from a model of *Stat1* immunodeficiency that permits MNV CR6, a non-hepatotropic enteric virus, to disseminate to the liver, where impaired viral replication in myeloid lineage cells leads to acute hepatic necroinflammation. Our data also suggest that non-hematopoietic (e.g. hepatocyte) cell-intrinsic STAT1 expression is a critical mediator of tissue resilience in the context of uncontrolled hepatic viral infection. Supporting this, when *Stat1* deficiency was restricted to immune populations, BM chimeras with STAT1-competent host cells recovered from acute infection-induced morbidity. In this context, persistent, subclinical hepatic CR6 infection was associated with ongoing myeloid cell dysregulation and fibrosis. Together, this model demonstrates that STAT1 functions in multiple cell types are necessary to prevent viral-induced hepatic damage. In the acute setting, STAT1 expression in infected cells is essential for effective viral containment and also promotes resilience of uninfected cells, whereas in the context of persistent infection, STAT1 inhibits AAMΦ differentiation and secondary fibrosis.

Conditional ablation of CD300lf in leukocyte populations has suggested that STAT1 plays an important role in restricting the cellular tropism of MNV CW3: in immunocompetent hosts, CD300lf deletion from LysM-expressing cells (primarily monocytes and MΦ), but not from neutrophils, DCs, or B cells, led to reduced CW3 viral burdens^36^. However, STAT1 deficiency broadened the cellular tropism for productive CW3 infection^36^. Consistent with these findings, our data demonstrate that STAT1 plays a similar role in restricting CR6 tropism. BM chimeras demonstrated that STAT1-deficient leukocytes, rather than non-hematopoietic epithelial cells or hepatocytes, are the primary source of CR6 replication. CR6 infection was detected in multiple myeloid lineage cells, including neutrophils, monocytes, MΦ, and DCs in the livers of *Stat1*^KO^ but not STAT1-sufficient mice. Although the relative contribution of unique myeloid cell lineages in propagating CR6-induced was not investigated, we have previously shown that neutrophils are dispensable for CR6-induced morbidity and mortality in *Stat1*^KO^ mice. Taken together with results demonstrating broad permissiveness for CW3 infection in *Stat1*^KO^ mice, the data suggest that coordinated dysregulation across multiple myeloid lineages, rather than individual cell types, could account for CR6 dissemination and tissue pathology.

Consistent with previous findings demonstrating that MNV CW3 persistence relies on inflammation-induced recruitment of a steady stream of myeloid cells susceptible to infection and lytic cell death^67^, CR6 infection in *Stat1*^KO^ mice is characterized by rapid MΦ recruitment into the liver. However, the initial source of hepatic infection that leads to the emergence of hepatic necroinflammatory lesions remains unresolved. It is possible that monocytes recruited from the circulation into the liver arrive already harboring infection. Alternatively, liver-resident Kupffer cells (KCs) may represent the first wave of CR6-susceptible myeloid cells. Independent of the inciting infection event, *Stat1*-deficiency in the immune compartment is both necessary and sufficient to establish a myeloid-rich hepatic environment that permits sustained CR6 infection.

Transcriptional analysis suggests additional consequences of CR6 infection in STAT1-deficient myeloid cells, including dysregulated activation and polarization. *In vivo*, uncontrolled lytic viral infection in *Stat1*^KO^ mice is associated with tissue damage and increased expression of myeloid inflammatory mediators at day 5 pi. Coincident with this, genes for alarmin cytokines and receptors (e.g. *il33*, *Il1rl1* (ST2)), and downstream effectors and receptors associated with type 2 immune-mediated tissue repair processes, including MΦ alternative activation (*chil3* (Ym1), *retnla* (RELMα)), are elevated in CR6-infected *Stat1*^KO^ mice. Because transcriptome analysis was performed on whole liver tissue, we cannot determine whether individual myeloid cells express genes encoding for effectors of both classically and alternatively activated MΦ, or if the signature results from mixed populations of classically activated CR6-infected cells and AAMΦ responding to tissue damage caused by lytic cell death and inflammation.

Our results also implicate a critical role for non-hematopoietic-derived cells (e.g. KCs, hepatocytes, hepatic stellate cells, liver sinusoidal endothelial cells) in the outcome of MNV CR6 infection. Transcriptional analysis of bulk liver tissue isolated from CR6-infected *Stat1*^KO^ mice indicated diminished expression of genes involved in aerobic respiration and xenobiotic metabolism. Hepatocytes make up the bulk of the liver, and they are critical regulators of organismal metabolism^68^. The high mitochondrial density in hepatocytes facilitates ATP generation via aerobic respiration, supporting essential processes of glucose homeostasis, urea genesis/ammonia detoxification, protein synthesis, and lipid storage. Within the xenobiotic metabolism category, genes in the cytochrome P450 (*cyp*), UDP-glucosyl transferase (*ugt)*, and glutathione-S-transferase (*gst*) families were down-regulated, collectively suggesting CR6 infection impairs liver detoxification and may lead to increased oxidative stress in *Stat1*^KO^ mice^69^. Whether the downregulation of genes and the associated metabolic processes are due to necroinflammatory hepatocyte loss or transcriptional reprogramming in viable hepatocytes, these disruptions likely contribute to local inflammation, systemic organ dysfunction, and mortality.

Results from mixed bone marrow chimeras support the hypothesis that STAT1-deficient hepatocytes or other non-hematopoietic-derived cell types contribute to CR6 pathogenesis. Despite harboring similar hepatic viral loads at day 7 pi, the extent of tissue inflammation and damage was more severe in CR6-susceptible *Stat1*^KO^→*Stat1*^KO^ chimeras vs resilient *Stat1*^KO^ →WT counterparts. Together, these data suggest that uncontrolled viral replication is not sufficient to account for disease outcome and implicate STAT1-deficient hepatocytes (or other hepatic cell types) as key drivers of acute tissue injury. It is possible that STAT1-competent hepatocytes are more resistant to lytic cell death induced by inflammatory or cytotoxic mediators produced by CR6-infected *Stat1*^KO^ myeloid cells. Because a STAT1-competent leukocyte compartment prevented (or effectively cleared) hepatic infection in WT→*Stat1*^KO^ chimeras, this hypothesis could not be directly examined *in vivo*.

Following resolution of acute CR6-induced weight loss, *Stat1*^KO^→WT chimeras appeared clinically healthy but harbored persistent CR6 infection at day 16 pi. This was associated with features of AAMΦ (e.g. *chil3* expression) and widespread hepatic fibrosis. Notably, in a model of pancreatic cancer, Ym1-expressing AAMΦ were identified as the primary source of fibrogenic TGFβ1^70^. These data align with reports implicating STAT1 as a negative regulator of both AAMΦ and liver fibrosis^63,71,72^. Currently, our understanding of STAT1 in liver fibrosis is largely based on its role in hepatocyte or hepatic stellate cell function^73–76^. Our data adds to this knowledge by demonstrating that STAT1-deficiency in the hematopoietic compartment is sufficient to drive fibrosis. Further analysis will be required to mechanistically dissect how persistent viral infection of *Stat1*^KO^ myeloid cells leads to activation and differentiation of hepatic stellate cells and/or portal fibroblasts into fibrogenic collagen-producing myofibroblasts. Additional analysis of the contribution of BM-derived fibrocytes to CR6-induced hepatic fibrosis in *Stat1*^KO^→WT BM chimeras may be warranted.

Collectively, the alignment of data from this mouse model of NHV pathology with relevant clinical findings suggest its utility for further analysis. In children born with STAT1 LOF mutations, hematopoietic stem cell transplant can mitigate the consequences of intracellular pathogen infections^12^. Consistent with this, WT BM reconstitution effectively restricted CR6 replication in hematopoietic cells and protected recipient *Stat1*^KO^ mice from hepatic infection and disease. Unfortunately, stem cell transplant does not restore STAT1 function in non-hematopoietic cell types and are not universally successful. Clinical reports indicated that 2 of 5 STAT1 LOF patients succumbed to post-transplant infections with EBV or CMV, viruses with non-hematopoietic cell tropism. Our data suggests that in addition to impaired antiviral immunity, STAT1-deficiency may render cells more susceptible to cellular stress and contribute to organ dysfunction or failure. Finally, the fibrosis associated with persistent hepatic CR6 infection in *Stat1*^KO^→WT chimeras may be applicable to settings of chronic HBV reactivation, or post-operative organ transplant patients recovering from acute NHV who should be monitored for post-viral sequelae, even after clinical symptoms subside.

## Methods

### Mouse strains and housing conditions

Mice were housed at the University of British Columbia (UBC) specific pathogen-free Centre for Disease Modeling. MNV-free C57BL/6J (000664), *Stat1*^KO^ (B6.129S(Cg)-*Stat1^tm1Dlv^/J,* 012606), and BoyJ (B6.SJL-*Ptprc^a^ Pepc^b^*, 002014) mice were sourced from Jackson Laboratories and bred in-house. Colonies were refreshed with new Jax-sourced C57BL/6J or BoyJ animals after 5 generations.

Multiple approaches were used to control for age, microbiome, and other environmental effects, including the use of littermates whenever possible, performing replicate experiments with litters from multiple dams, and co-housing mice of mixed genotypes prior to and throughout experiments. We have previously shown that *Stat1*^Het^ animals are suitable STAT1-sufficient controls^51^. Thus, *Stat1*^+/-^ x *Stat1*^-/-^ breeders were used to generate experimental *Stat1*^Het^ and *Stat1*^KO^ littermates. For larger experiments, age-matched mice from multiple litters were included. Matched litters from brother–sister harems were used in transcriptomic and bone marrow chimera experiments to control for genetic and environmental variability. Animals were sex-matched within experiments and replicates were stratified by sex to assess sex-specific differences (none detected, not shown). Experimental mice were used between 6-12 weeks of age and were aged-matched within individual experiments.

All experiments and breeding were performed in accordance with UBC Animal Care Committee and Biosafety Committee-approved protocols. Mice were housed in ventilated Ehret cages prepared with BetaChip bedding and had *ad libitum* access to irradiated PicoLab Diet 5053 and reverse osmosis/chlorinated (2-3 ppm)-purified water. Housing rooms were maintained on a 14/10-hour light/dark cycle with temperature and humidity ranges of 20-22°C and 40-70%, respectively. Sentinel mice housed in experimental rooms were maintained on dirty bedding and nesting material and were tested on a quarterly basis for presence of mites (*Myocoptes, Radford/Myobia)*, pinworm (*Aspiculuris tetaptera, Syphacia obvelata*), fungi (*Encephalitozooan cuniculi*), bacteria (*Helicobacter* spp., *Clostridium piliforme, Mycoplasma pulmonis,* CAR Bacillus), and viruses (Ectromelia, EDIM/Rotavirus, MHV, MNV, MPV, MVM, LCMV, MAV1/2, MCMV, Polyoma, PVM, REO3, Sendai and TMEV).

### MNV preparation, infections and quantification

MNV CR6 was grown and isolated from RAW264.7 cells as previously described^44,77^. Mice were infected i.v. with 2x10^5^ plaque forming units (pfu) or 2.9x10^5^ fifty-percent tissue culture infectious dose (TCID_50_) of MNV CR6 diluted in sterile PBS. Naive mice were sham-infected with sterile PBS. Mice were monitored every 24-48 hours and euthanized at humane endpoint (loss of ≥20% of initial body weight). In rare cases, true infection-induced mortality occurred in *Stat1*^KO^ mice that did not meet the criteria of humane endpoint.

MNV genome copies were quantified in tissue samples that had been collected into RNAlater and stored at -80°C until use. Thawed tissues were removed to clean tubes containing PureLink RNA Mini (ThermoFisher) lysis buffer, disrupted with bead beating in a TissueLyser II (Qiagen), and RNA extracted from cleared supernatants according to manufacturer instructions. cDNA was generated from RNA using Superscript III Reverse Transcriptase primed with random hexamers (ThermoFisher). MNV genome copies were amplified in PerfeCTa qPCR FastMix II, Low ROX (VWR) with primers MNV F4972–(5’ - CAC GCC ACC GAT CTG TT– TG - 3’), R5064–(5’ - GCG CTG CGC CAT CA– TC - 3’) and probe 5001-5015 (–FAM - CGC TTT GGA ACA–ATG -MGBNFQ) on a Quantstudio 5. Absolute numbers of MNV genome copies were interpolated from a standard curve and normalized to tissue weight.

Infectious virus was titrated by TCID_50_ assay in 96-well plates. Tissue was weighed, bead-homogenized (30Hz for 6 minutes, Qiagen TissueLyser II), and serially diluted in serum-free media (neat to 10^-4^) before being overlaid on RAW264.7 cells (8 replicates per dilution, per sample). Plates were rocked at RT for 1 hour, inoculum aspirated and DMEM CTCM added (containing 10% FBS, 50 U/ml penicillin/streptomycin, 25 mM HEPES, and 55 µM β-Mercaptoethanol). Cells were incubated (37°C and 5% CO_2_) for 72 hours, then stained with crystal violet and visually scored for cytopathic effect (binary scoring, CPE visible or not). TCID_50_ was calculated by the Reed-Muench method and normalized to sample weight^78^.

### Hepatic leukocyte recovery

Mice were euthanized by CO_2_ inhalation and transcardially perfused with at least 10 mL cold PBS, until visible liver lightening. The left lateral lobe of the liver of PBS-perfused animals was collected for isolation of hepatic leukocytes. Liver sections were minced with fine scissors and enzymatically digested (10 ml DMEM + 2% NCS with 2.5mg/mL collagenase D (Roche Diagnotics), 50 µg/mL DNAse I (Sigma)) at 37°C with 180 rpm shaking for 30 minutes. Samples were passed through a 70 µm nylon mesh filter and centrifuged at 50 x g for 3 minutes to at RT to pellet hepatocytes. Cells recovered from the supernatant were subjected to ACK hemolysis. Cells were resuspended in 40% Percoll in DMEM and centrifuged at 600 x g (minimum acceleration and braking) for 20 minutes at RT. Lipid and contaminant-containing supernatant was discarded. The leukocyte-enriched cell pellet was washed and prepared for flow cytometric analysis by resuspension in FACS buffer (PBS, 2% newborn calf serum (NCS), 2 mM EDTA).

### Flow cytometry

Cells were stained with LIVE/DEAD Fixable Aqua Dead Cell Stain (Invitrogen) in PBS prior to blocking and antibody staining in FACS buffer. Anti-mouse CD16/CD32 antibody clone 2.4G2 was included at 2.5 µg/mL in staining cocktails with the antibodies listed in Table 1. Flow cytometry was performed on a Cytoflex (Beckman-Coulter) flow cytometer, and results were analysed using FlowJo (version 10). Cell sorting was performed on a Cytoflex SRT (Beckman-Coulter).

**Table 1.**
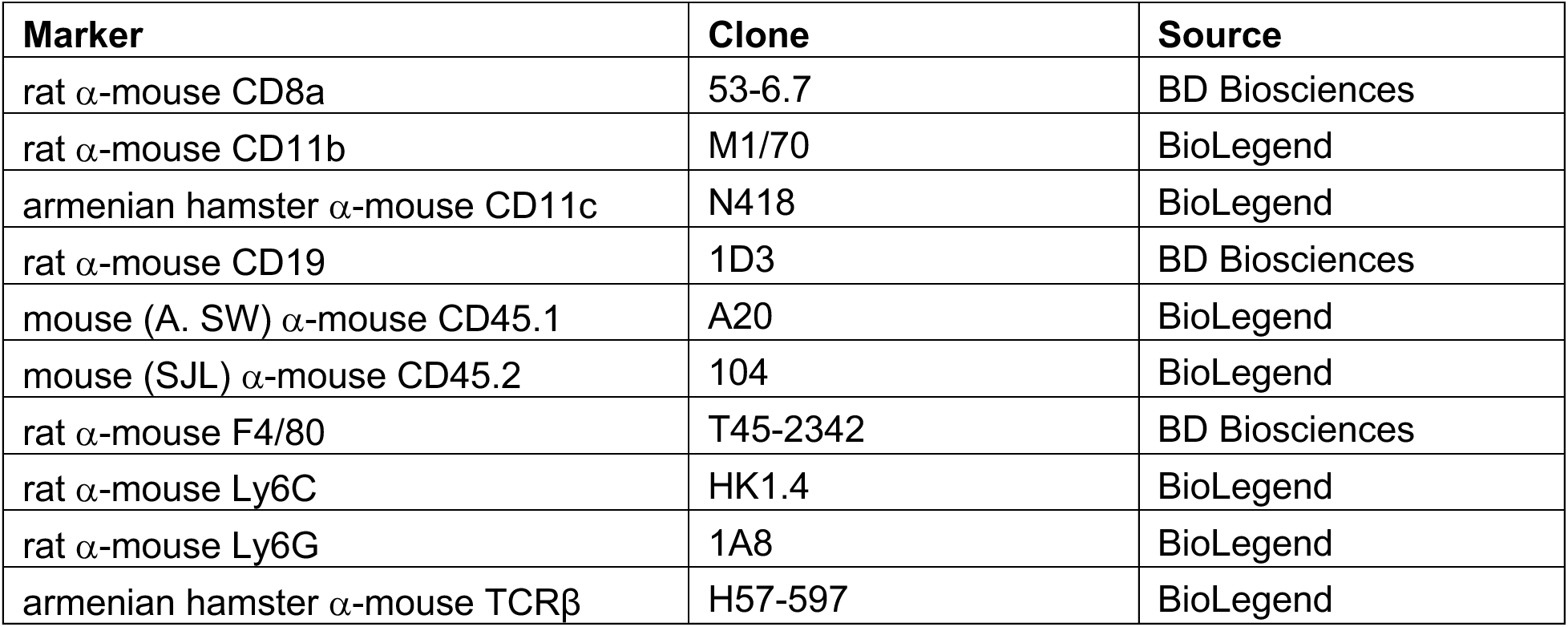
Antibodies used for flow cytometry.

### Histology and Immunofluorescence

Perfused tissues were fixed in 4% paraformaldehyde and embedded in paraffin. Hematoxylin & eosin and Masson’s trichrome staining was performed on 5 <u>µ</u>m sections of liver at the UBC Histochemistry Service Laboratory. Automated quantification of diseased area in liver samples was performed on tiled images of entire liver sections (ImageJ). Briefly, colour deconvolution was performed and thresholding for diseased area applied (hue, saturation, and brightness, which were first determined manually for each experiment). Diseased area was calculated as a percent thresholded area of the whole liver image.

NS1 staining was performed as previously described^46,53,79^. Formalin-fixed paraffin-embedded (FFPE) tissue sections (7 µm) were deparaffinized by heating for 20 minutes at 65°C followed by a xylene rinse and subsequently rehydrated by sequential rinses in increasingly dilute ethanol solutions. Samples were incubated for 15 minutes at 110°C in Dako Target Retrieval Solution (Agilent) and an NxGen Decloaking Chamber (Biocare Medical) for antigen retrieval. Slides were blocked in PBS with 1% NCS and 2% mouse serum prior to staining. Primary monoclonal antibody rabbit α-mouse CD11b (Abcam ab133357) was diluted 1/500 and/or Armenian hamster α-MNV NS1 clone 11G10^79^ was diluted 1/1000 in a staining buffer consisting of PBS with 1% newborn calf serum and added to samples overnight at 4°C. Secondary antibody donkey α-rabbit AlexaFluor680 (Invitrogen A10043) was diluted 1/400, goat α-rabbit AlexaFluor488 (Life Technologies A11034) was diluted 1/500, or goat α-hamster AlexaFluor647 (BioLegend 405510) was diluted 1/500 in staining buffer and added to samples for 2 hours at 4°C. Tissues were counterstained with DAPI (4′,6-diamidino-2-phenylindole) and mounted using VectaMount mounting medium (Vector Laboratories). Histology and immunofluorescent images were acquired with Zeiss Axio Observer 7 and an AxioCam 105 microscope camera.

### RNAscope

Slides with 7 mm formalin-fixed paraffin-embedded tissue sections were treated with RNAscope Multiplex Fluorescent Detection Kit v2 (Advanced Cell Diagnostics, Newark, CA) for in situ hybridization with MNV CR6 probes^53,80^. Briefly, tissues were hybridized with custom-designed probes specific for MNV CR6 RNA for 2 h at 40°C, and the signal was detected using fluorescent Opal dyes (Akoya Biosciences). Opal 520 was used for MNV CR6 probes. Tissues were counterstained with DAPI (4′,6-diamidino-2-phenylindole) and mounted using VectaMount mounting medium (Vector Laboratories). Images for FISH were acquired using a Nikon A1R confocal microscope, and images were captured using NIS Elements (Nikon) software.

### RNA differential gene expression analysis

Fresh perfused liver sections from the left lateral lobe were collected and snap-frozen in TRIzol (ThermoFisher). Thawed tissues were lysed by bead beating in a QIAgen TissueLyser II and RNA was isolated from cleared supernatants according to manufacturer’s instructions. The resulting RNA was assessed on an Agilent 2100 Bioanalyzer and samples with RNA integrity numbers > 8 were prepped following the standard protocol for Illumina Stranded mRNA prep (Illumina). Sequencing was performed on the Illumina NextSeq2000 with Paired End 59bp × 59bp reads. FASTQ files were mapped and aligned using the Salmon Aligner v1.10.1. Reads were mapped to an index created using the GRCm38 mouse transcriptome assembly with decoy sequences generated using SalmonTools. Mapping and alignment was performed on the UBC Sockeye Advanced Research Computer. Differential gene expression analysis was performed on mapped and aligned files output from Salmon in R using DESeq2 and ClusterProfiler. Gene set enrichment analysis with ClusterProfiler compared differential gene expression data to publicly available datasets (GSE9892^57^, GSE216278^58^, GSE214778^59^, GSE222576^60^, GSE138263^62^) and all mouse Gene Ontology terms. For human datasets (GSE62037^61^), gene identifiers were first converted to homologous mouse IDs using BioMaRt and the JAX homology database^81^. Heatmaps were generated with the pheatmap function, with genes hierarchically clustered by Ward’s method on Euclidean distances. Heatmap data represent mean-normalized expression values. Throughout RNA analyses, criteria for significance were log_2_ fold change > 2 or < -2, p_adj_ < 0.05, and base mean > 50, with p values adjusted for multiple comparisons using FDR cutoff of 0.05 (5%).

### BMDM generation and infection

Bone marrow was isolated from the tibias and femurs of *Stat1*^KO^ and *Stat1*^Het^ co-housed littermates that were less than 18 weeks of age. Resulting cells were counted and plated in tissue culture treated six well plates at 10^6^ cells per well. BMDMs were cultured in Dulbecco’s Modified Eagle Medium complete total cell media (DMEM CTCM, containing 10% FBS, 50 U/ml penicillin/streptomycin, 25 mM HEPES, and 55 µM β-Mercaptoethanol) and supplemented with macrophage colony stimulating factor (20 ng/well, Gibco) on days 0, 3, and 6. BMDMs were incubated at 37°C and 5% CO_2_, and used experimentally at day 7. Sterile serum-free DMEM or 5 x 10^5^ pfu CR6 was added to cultures following a wash with serum-free media and were rocked for one hour at room temperature. Following infection, supernatant was discarded, DMEM CTCM was added, and cells were incubated for 24 hours before collection in RNAlater.

### RNA extraction and qRT-PCR

RNA was extracted from cells lysed using a TissueLyser II (Qiagen) at 30Hz for 6 minutes with the PureLink RNA Mini Kit (Thermo) and reverse transcribed (SuperScript III Reverse Transcriptase, Thermo). Subsequent cDNA was amplified (QuantStudio 5, Thermo) with specified primers (Table 2) and either a TaqMan probe for MNV quantification in triplicate (against a standard curve, as described above) or SYBR green dye for gene expression in duplicate. For gene expression qPCR, fold change was determined through the ΔΔC_t_ method and internally controlled with *hprt*.

**Table 2.**
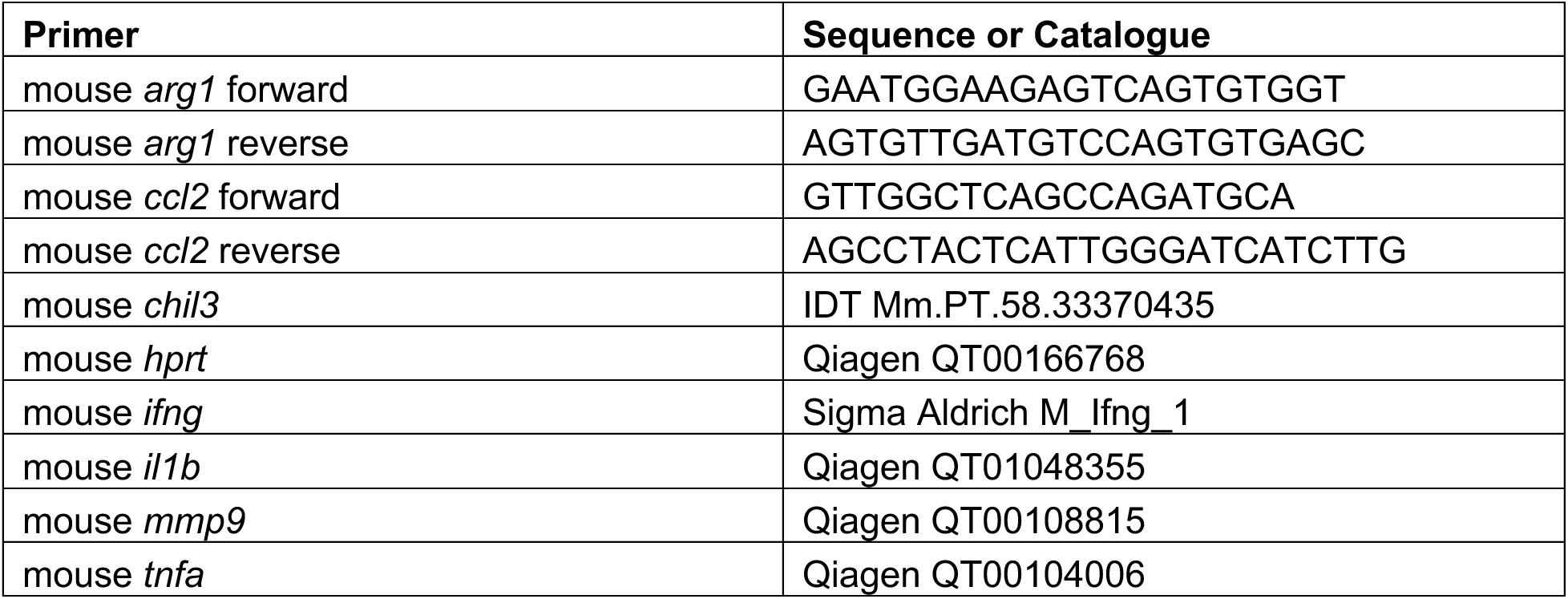
Primers for gene expression qPCR.

### Bone marrow chimeras

CD45.2^+^ *Stat1*^KO^ and CD45.1^+^ BoyJ mice were irradiated with 650 rads twice, 4 hours apart in groups of 3-4. Bone marrow was isolated from naive CD45.2^+^ *Stat1*^KO^ and CD45.1^+^ BoyJ mice 24 hours post-irradiation and ACK lysed to remove erythrocytes. Irradiated mice were i.v. injected with 10^6^ bone marrow cells diluted in sterile PBS. Mice were supplemented with 2 mg/ml Neomycin in drinking water for 14 days, DietGel was provided for 10 days, and mice were allowed to reconstitute for 6 weeks prior to MNV infection. Chimeric reconstitution was confirmed by flow cytometry.

### Statistical Analyses

Statistical testing was performed in GraphPad Prism (version 10), and appropriate tests were selected as indicated in figure legends. On all graphs, bars indicate median values, and points represent biological replicates. Values in flow cytometry plots indicate average value for indicated group ± group standard deviation. Statistically significant differences are shown in figures as follows: *, p < 0.05; **, p < 0.01; ***, p < 0.001; ****, p < 0.0001; and ns, not significant.

## Data Transparency

This study did not generate new unique reagents. RNAseq data is available from NCBI’s Gene Expression Omnibus and are accessible through GEO Series accession number GSE326527. Materials or other additional information required to reanalyze the data reported in this paper will be made available by the lead contact upon request.

## Acknowledgements

Research in the Osborne lab takes place on the traditional, ancestral, and unceded territory of the xwməqkwəýəm (Musqueam) First Nation. We are grateful for technical expertise provided by UBC core facilities, including Andy Johnson and Justin Wong at the Life Sciences Institute ubcFLOW core, Tara Stach and Yvonne Chung of the UBC SBME-seq core, and the Histochemistry Service Laboratory at UBC’s Center for Comparative Medicine. Our work would not be possible without the animal care staff at the Centre for Disease Modeling. Many thanks to Dr Phil Domeier for thoughtful feedback on this manuscript.

